# Novel phosphorylation of Rab29 that regulates its localization and lysosomal stress response in concert with LRRK2

**DOI:** 10.1101/2022.09.26.509472

**Authors:** Tadayuki Komori, Tomoki Kuwahara, Tetta Fujimoto, Maria Sakurai, Takeshi Iwatsubo

**Affiliations:** Department of Neuropathology, Graduate School of Medicine, The University of Tokyo, Tokyo, Japan

## Abstract

Rab proteins are small GTPases that regulate a myriad of intracellular membrane trafficking events. Rab29 is one of the Rab proteins phosphorylated by leucine-rich repeat kinase 2 (LRRK2), a Parkinson’s disease-associated kinase. Recent studies suggest that Rab29 regulates LRRK2, whereas the mechanism by which Rab29 is regulated remained unclear. Here we report a novel phosphorylation in Rab29 that is not mediated by LRRK2 and occurs under lysosomal overload stress. Mass spectrometry analysis identified the phosphorylation site of Rab29 as Ser185, and cellular expression studies of phosphomimetic mutants of Rab29 at Ser185 unveiled the involvement of this phosphorylation in counteracting lysosomal enlargement. PKCα was deemed to be responsible for this phosphorylation and control the lysosomal localization of Rab29 in concert with LRRK2. These results implicate PKCα in the lysosomal stress response pathway comprised of Rab29 and LRRK2, and further underscore the importance of this pathway in the mechanisms underlying lysosomal homeostasis.

## Introduction

The Rab small GTPases are the largest protein family in the Ras superfamily, with approximately 60 proteins identified. These Rab GTPases (Rabs) are often called the master regulators of membrane traffic, modulating the biogenesis, trafficking, tethering, or fusion of membranes through various effector proteins. Rab function requires two important modes of regulation. The first is the C-terminal prenylation of Rabs that happens after synthesis and is responsible for its membrane association. The second is its guanine nucleotide binding state, which is controlled by guanine nucleotide exchange factors (GEFs) and GTPase activating proteins (GAPs). GEFs exchange the GDP bound to Rabs to GTP to make Rabs active; GAPs promote hydrolysis of GTP bound to Rabs to make them inactive. Both GEFs and GAPs have a high specificity for each Rab, but activation only occurs on membranes^1–4^.

In addition to activation by altering nucleotide binding states, there are several other means of regulating Rab activity. Among them is phosphorylation, which was associated with altered subcellular localization in early studies^5,6^. This post-translational modification has gained much attention after the discovery of a subset of Rabs being phosphorylated by leucine rich-repeat kinase 2 (LRRK2), a Parkinson’s disease (PD) causative protein^4,7,8^. Recently, it has been reported that more than 40 Rab and related proteins are phosphorylated upon activation of a platelet collagen receptor GPVI^9^, further indicating the importance of Rab regulation by phosphorylation.

Rab29 is one of the Rab proteins identified to be phosphorylated by LRRK2^10–12^. *RAB29* is thought to be a risk factor of PD, encoded in a susceptibility locus *PARK16* ^13–15^, further implicating the link with LRRK2. The consensus of the main function of Rab29 lies in the regulation of LRRK2^16^, with multiple reports observing the activation of LRRK2^17^, recruitment of LRRK2 to the Golgi upon overexpression^12,18^ or to lysosomes upon lysosomal overload^19^. Other intracellular functions of Rab29 include Golgi-associated trafficking^20,21^, AP3-associated protein trafficking^22^, maintenance of lysosomal homeostasis^19^, and Golgi-independent endolysosomal trafficking^23^, most of which are functions related to LRRK2. Recent analyses of the direct interaction of these two proteins, with the help of improved structure analysis techniques, have revealed the position and surface of their interaction^24–26^, further lighting the way to analyze the Rab29-LRRK2 axis.

However, apart from the regulation of LRRK2 by Rab29, the upstream mechanism regulating Rab29 itself remains largely elusive. We have previously reported that LRRK2-mediated phosphorylation of Rab29 at Ser72 regulates Golgi morphology^11^. There are also reports that the neighboring Thr71 can also be phosphorylated together^10,12^, which suggest the importance of Rab29 phosphorylation at these sites in steady-state conditions. Under stressed conditions that cause lysosomal overload, Rab29 is translocated from Golgi to the lysosomal surface and regulates the morphology of lysosomes^19^, whereas the regulatory mechanisms or factors of Rab29 under these conditions are yet to be known. Here we report a novel phosphorylation of Rab29 at Ser185 that, together with the phosphorylation by LRRK2, regulates the lysosomal stress response, shedding light on further insights into the pathophysiological functions of Rab29.

## Results

### Lysosomal stress leads to LRRK2-independent phosphorylation of Rab29

We have previously reported that overloading lysosomes causes sequential recruitment of Rab29 and LRRK2 onto enlarged lysosomes^19^ and that this recruitment is brought about by various lysosomotropic agents that cause lysosomal overload^27^.

Therefore, we assumed that there could be a mechanism underlying Rab29 localization to lysosomes under conditions causing lysosomal overload. Chloroquine (CQ), a lysosomotropic agent, was used to elicit lysosomal overload stress in HEK293 cells overexpressing Rab29. We found that Rab29 was phosphorylated upon exposure to CQ, and that the phosphorylation increased over time **(Fig. 1A)**. This phosphorylation was also observed in endogenous Rab29 in HEK293 or RAW264.7 cells **(Fig. 1B)**.

**Figure 1.**
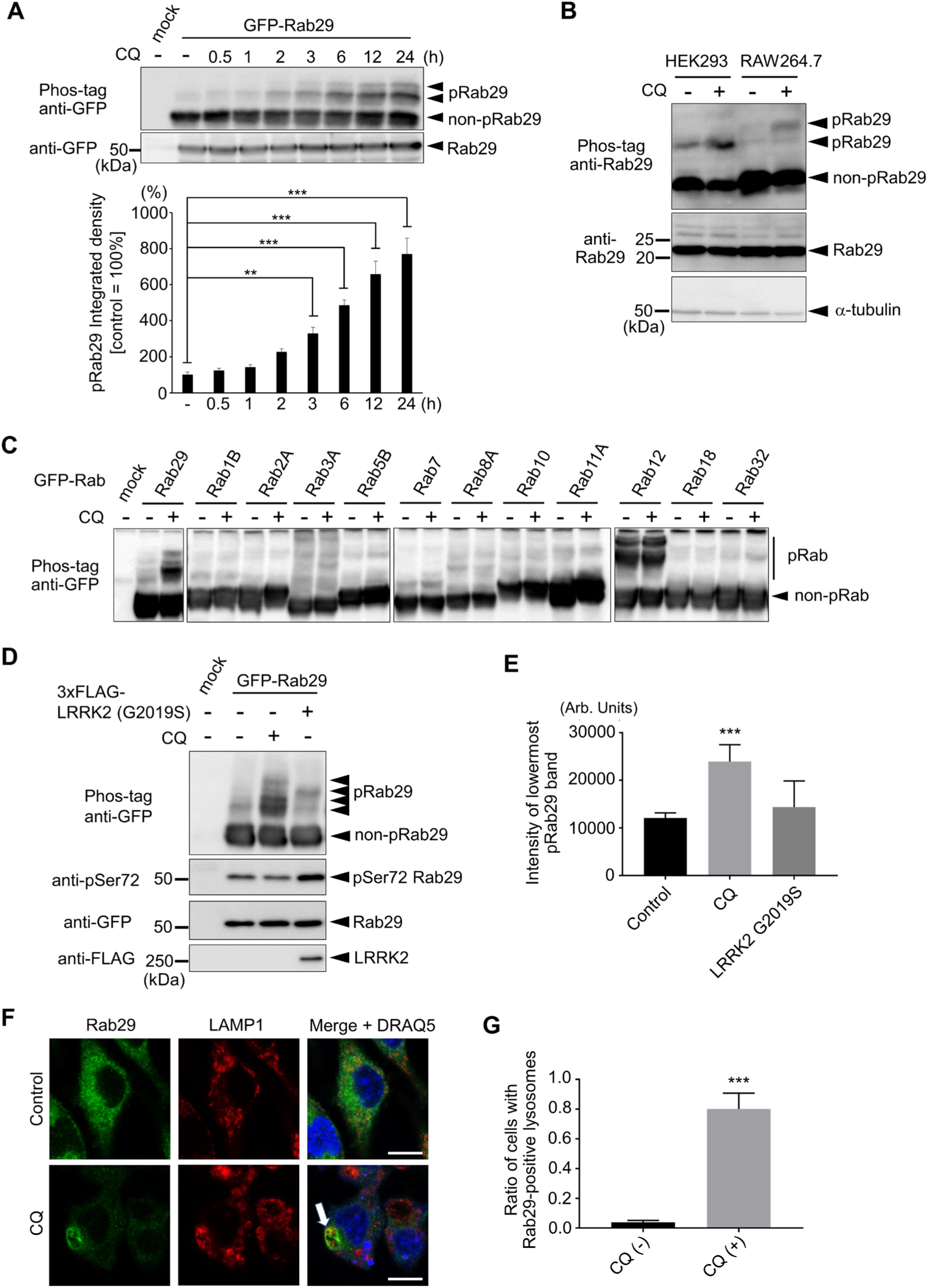
Lysosomal stress leads to LRRK2-independent phosphorylation of Rab29. **(A)** Phosphorylation of Rab29 by CQ treatment in a time-dependent manner. pRab29 andnon-pRab29 indicate phosphorylated and nonphosphorylated Rab29, respectively. One-way ANOVA followed by Dunnett’s test against the control. **: *p* < 0.01, ***: *p* < 0.001. **(B)** Phosphorylation of endogenous Rab29 in RAW264.7 and HEK293 cell lines. **(C)** Phosphorylation of a set of Rab proteins with or without CQ. **(D)** Comparison of Rab29 phosphorylation by CQ and LRRK2. **(E)** Quantification of the intensity of the lowermost phosphorylated band in D. One-way ANOVA followed by Dunnett’s test against the control. ***: *p* < 0.001. **(F)** Localization of endogenous Rab29 under CQ-treated conditions. The arrow indicates Rab29 colocalization with LAMP1, a lysosomal marker. Bars = 10 μm. **(G)** Quantification of lysosomal localization of Rab29, as shown in F. Student’s *t*-test against the control. ***: *p* < 0.001

To see whether this phenomenon is prevalent among other members of the Rab family, CQ was treated to HEK293 cells expressing various Rab proteins. Induction of phosphorylation was prominent only in Rab29, but was not observed in any other Rabs tested **(Fig. 1C)**, including Rab32, the closest Rab to Rab29 in the Rab subfamily^4^, and Rabs 3A, 8A, 10 and 12 that are well known substrates of LRRK2^8^. This difference raised the possibility that LRRK2 is not likely to be the kinase responsible for this Rab29 phosphorylation caused by CQ.

Next, we compared Rab29 phosphorylation by LRRK2 with that by CQ. Co-expression of LRRK2 resulted in phosphorylation as detected in a Phos-tag SDS-PAGE analysis, but its band intensities and patterns were different from those seen upon CQ treatment **(Fig. 1D, E)**. Furthermore, the signal for phospho-Ser72, a residue phosphorylated by LRRK2^11^, was unaltered in CQ-treated cells **(Fig. 1D)**, further supporting the LRRK2-independency of this phosphorylation.

Also in our previous study using RAW264.7 cells, Rab29 was found to mediate the translocation of endogenous LRRK2 to enlarged lysosomes under CQ exposure^19^. We therefore assumed that the localization of endogenous Rab29 could also change in the same manner as LRRK2. Immunocytochemical analysis revealed that CQ treatment caused the translocation of endogenous Rab29 to enlarged lysosomes, as observed with LRRK2 **(Fig. 1F, G)**.

### Phosphorylation of Rab29 occurs on the lysosomal surface

Since CQ exposure caused Rab29 phosphorylation and its translocation to enlarged lysosomes, we speculated that this phosphorylation could occur at the lysosomal membrane. To explore this possibility, we utilized a forced intracellular protein translocation system^28^, a drug-inducible method based on the heterodimerization of the FK506-binding protein (FKBP) and the FKBP-rapamycin binding domain (FRB) in the presence of rapamycin analogs **(Fig. 2A)**. FKBP-tagged Rab29 expressed in HEK293 cells was forced to localize at the lysosomal surface by co-expressing the lysosomal protein LAMP1-FRB followed by treating with AP21967, a rapamycin analog heterodimerizer. Similar to CQ treatment, phosphorylation of Rab29 increased over time upon treatment with AP21967 **(Fig. 2B)**.

**Figure 2.**
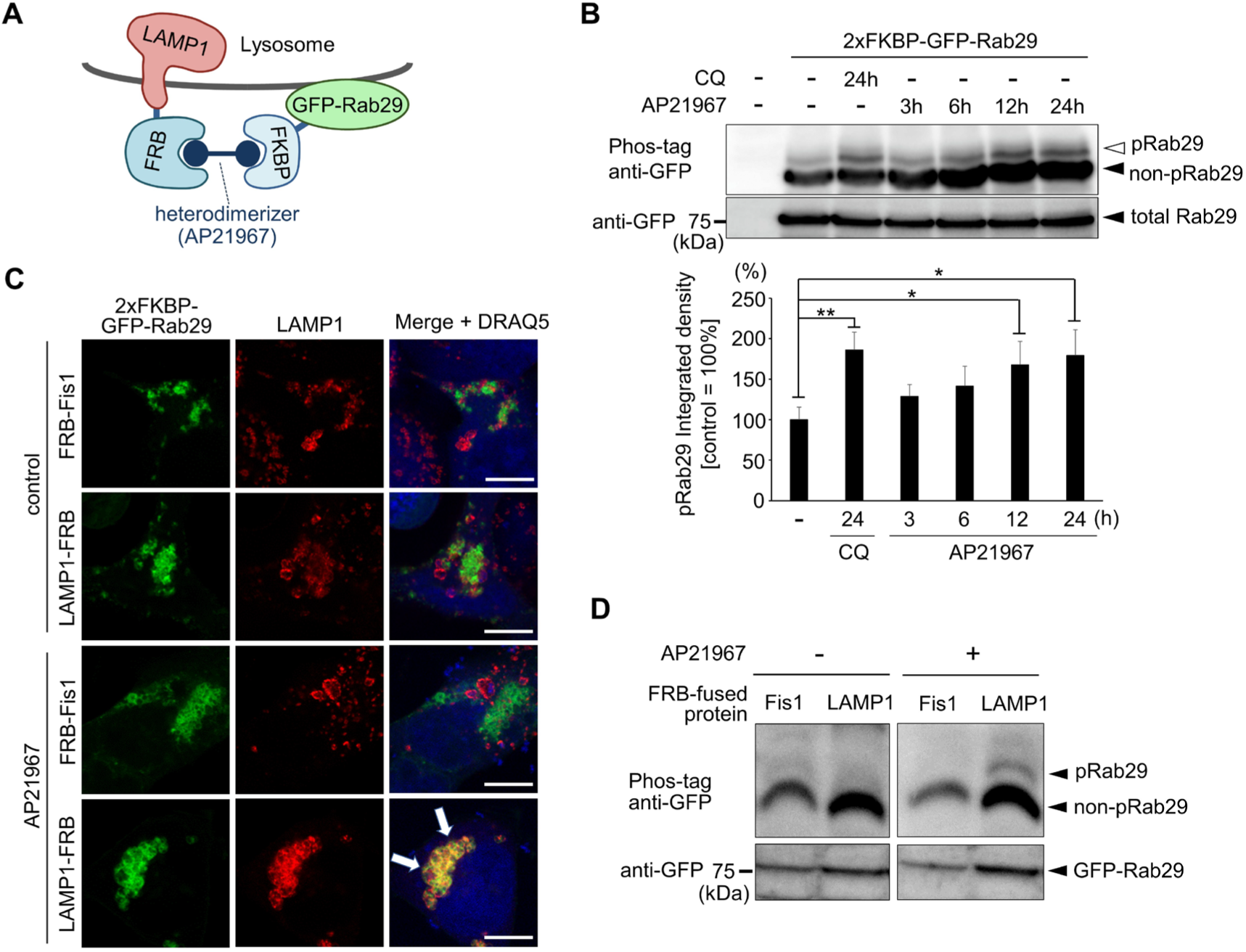
Phosphorylation of Rab29 occurs on the lysosomal surface. **(A)** A scheme of the forced localization technique used in this study. Upon treatment with the heterodimerizer AP21967, FKBP-bound Rab29 is anchored away to FRB-positive compartments in the cell. **(B)** Phosphorylation of FKBP-Rab29 over time upon lysosomal forced localization by AP21967 in HEK293 cells co-expressing LAMP1-FRB. One-way ANOVA followed by Dunnett’s test against the control. *: *p* < 0.05, **: *p* < 0.01. **(C)** Anchoring Rab29 towards mitochondria (Fis1) or lysosomes (LAMP1). Arrows indicate Rab29 colocalized with LAMP1. Bars = 10 μm. **(D)** Phosphorylation of Rab29 upon forced localization to lysosomes (LAMP1) but not to mitochondria (Fis1).

We confirmed that the intracellular localization of Rab29 changed to lysosomes when LAMP1-FRB was co-expressed and treated with AP21967 **(Fig. 2C)**. This lysosomal localization was not observed when mitochondrial protein FRB-Fis1 was co-expressed instead of LAMP1-FRB **(Fig. 2C)**. We found that the co-expression of LAMP1-FRB, but not FRB-Fis1, induced the phosphorylation of Rab29 **(Fig. 2D)**, indicating that the phosphorylation of Rab29 occurred on the lysosomal surface.

### Ser185 residue of Rab29 is phosphorylated under lysosomal stress

As we assumed that this phosphorylation is independent of LRRK2, we sought for the exact phosphorylation site of Rab29. After pulling down GFP-Rab29 from CQ-treated cells, phosphorylated Rab29 was separated by Phos-tag SDS-PAGE and analyzed by mass spectrometry, which identified the putative phosphorylation sites as Ser31 or Ser185 **(Fig. 3A, B)**. Ser31 is just before the switch 1 region and Ser185 is in the third complementarity determining region (CDR3)^7,29^, which is a motif that contributes to the specificity of its effector^30^. Both sites harbor a potential to regulate the activity of this GTPase.

**Figure 3.**
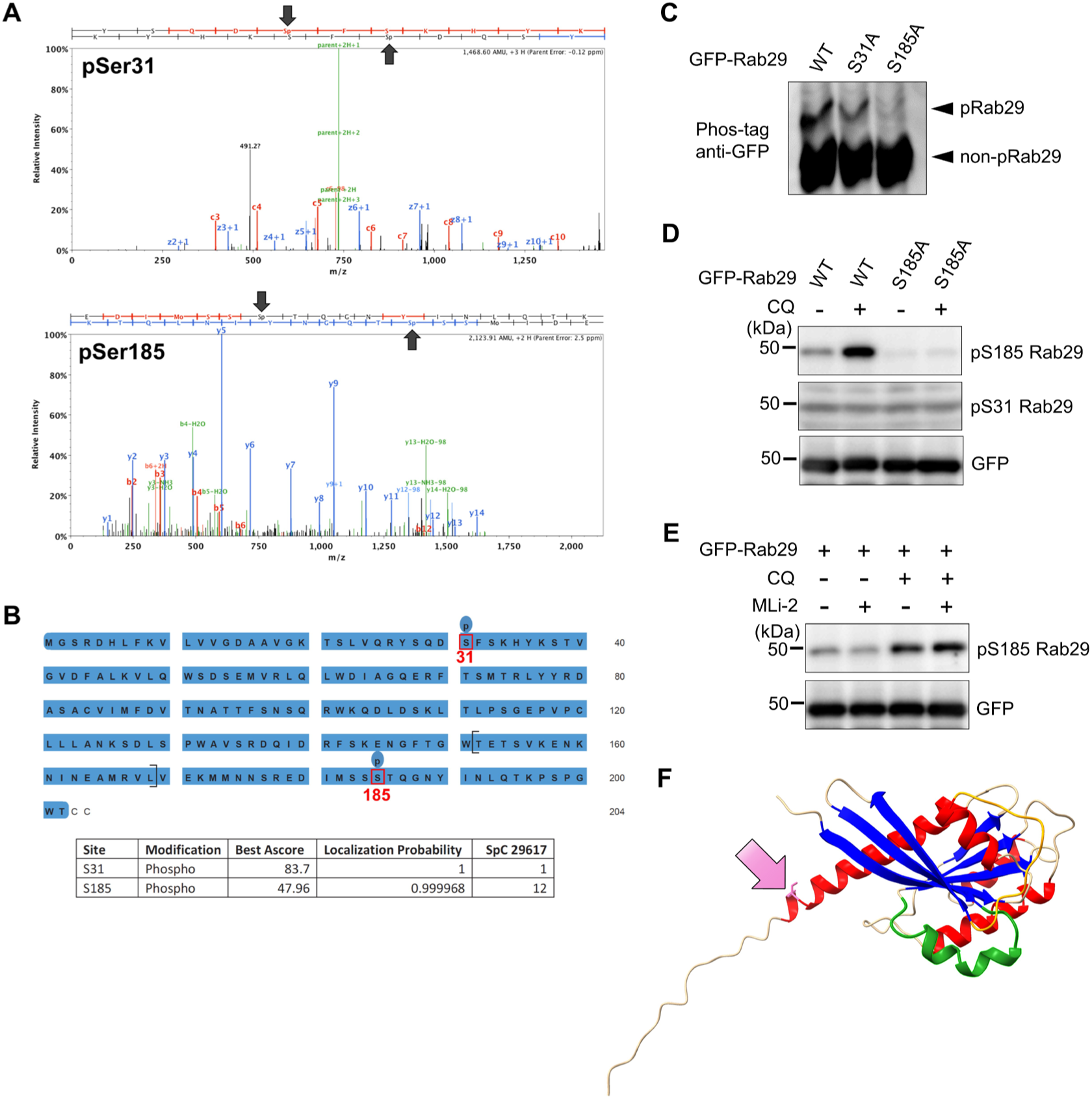
Ser185 residue of Rab29 is phosphorylated under lysosomal stress. **(A)** A representative spectrum of LC-MS/MS indicating phosphorylation at Ser31 or Ser185. **(B)** A sequence of Rab29 showing the putative phosphorylation sites Ser31 and Ser185. **(C)** Reduced phosphorylation of Rab29 by alanine substitution of Ser185, but not Ser31, under CQ treatment. **(D)** Confirmation of Ser185 phosphorylation by using phospho-specific antibodies. **(E)** No changes in CQ-induced phosphorylation of Rab29 by inhibition of LRRK2. **(F)** An AlphaFold2-generated structural image of Rab29. The pink residue pointed by a pink arrow is Ser185. The orange and green chains indicate Switch 1 and 2, respectively. α-helices are colored in red and β-sheets in blue.

To further determine the phosphorylation sites, alanine mutants Ser31 to Ala (S31A) or Ser185 to Ala (S185A) were overexpressed in HEK293 cells followed by treatment with CQ. The phos-tag SDS-PAGE revealed a major decrease of the phosphorylated Rab29 band in the S185A mutant, compared to either S31A or wild-type (WT) Rab29 **(Fig. 3C)**. Thus, the phosphorylation of Rab29 under CQ treatment was thought to occur at Ser185.

To further examine the Rab29 phosphorylation, we developed a phospho-Ser185 specific rabbit polyclonal antibody and analyzed Rab29-overexpressing cells. We found that the signal recognized by this antibody was elevated upon CQ exposure and was abolished in the S185A mutant **(Fig. 3D)**. Treatment with a LRRK2-specific kinase inhibitor, MLi-2, did not affect the band intensity of phospho-Ser185 **(Fig. 3E)**, confirming that this phosphorylation is indeed independent of LRRK2.

According to a structural prediction by AlphaFold2^31^, the Ser185 residue is located at the end of the last helix **(Fig. 3F)**. As this site lies in CDR3^7,29^ and phosphorylations in this region could alter binding with GEFs or GDIs^7^, it seemed reasonable to assume that this phosphorylation could somehow regulate the function of Rab29.

### Phosphomimetics of Ser185 alleviated CQ-induced lysosomal enlargement

We next sought to analyze the role of this phosphorylation in terms of lysosomal regulation. Phosphorylation of Ser185 residue was mimicked by substitution to aspartate or glutamate (S185D or S185E, respectively), or disabled by alanine substitution (S185A). These mutants were overexpressed in HEK293 cells and their effects on Rab29 localization or lysosome morphology were assessed. Under steady-state conditions, overexpression of these mutants did not alter their subcellular localization, and the morphology of lysosomes remained small and unaltered **(Fig. 4A)**. Upon CQ treatment, WT or S185A Rab29-overexpressing cells exhibited enlarged lysosomes as observed in previous studies, whereas S185D or S185E Rab29-overexpressing cells exhibited smaller lysosomes, similar to those observed in CQ-untreated samples **(Fig. 4B, C)**.

**Figure 4.**
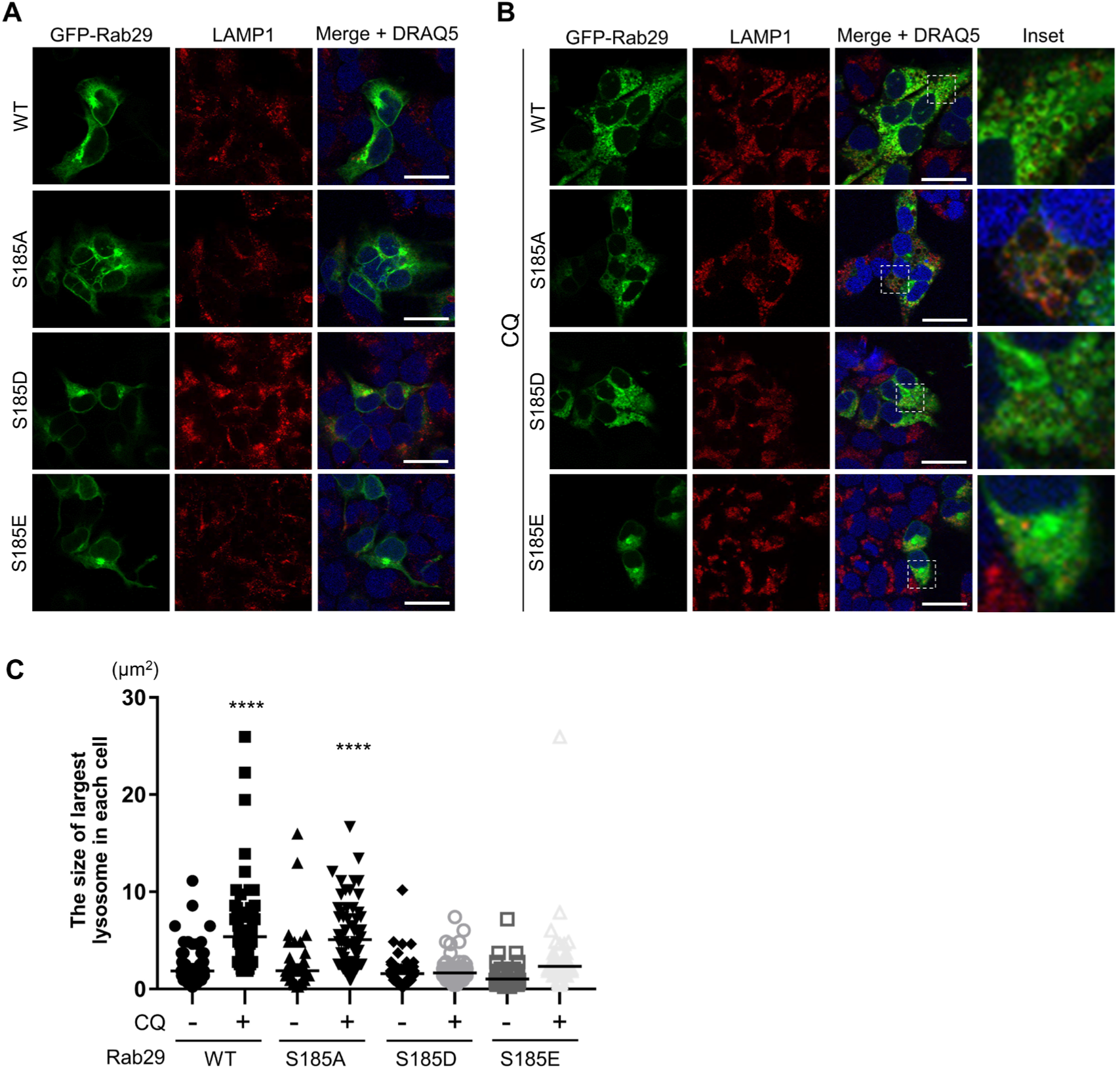
Phosphomimetics of Ser185 alleviate CQ-induced lysosomal enlargement. **A)** Lysosome morphology and Rab29 localization in HEK293 cells expressing Rab29 WT or Ser185 mutants at steady state. **(B)** Lysosome morphology and Rab29 localization in HEK293 cells expressing Rab29 WT or mutants upon CQ treatment. Insets of areas containing lysosomes are shown on the right. Bars = 10 μm. **(C)** Statistical analysis of lysosomal size in A and B. Each shape shows the area of the largest lysosome in each cell, obtained by elliptical approximation of each immunocytochemistry image. The mean is shown by a black horizontal bar in each sample. One-way ANOVA followed by Dunnett’s test against the control (wild type, no CQ) sample, ****: *p* < 0.0001.

LRRK2 inhibition or knockdown has been shown to increase lysosomal size when treated with CQ^19,27^. Also, several studies have reported that overexpression of Rab29 increases the LRRK2 kinase activity^17,27^. Therefore, we next examined whether any of these mutant Rab29 would alter the LRRK2 kinase activity. Using Rab10 phosphorylation at Thr73 as a readout^19,27,32^, LRRK2 kinase activity was measured and no significant changes were observed **(Supplementary Figure 1A, B)**. Also, no changes in LRRK2 binding to Rab29 mutants compared to WT Rab29 were observed **(Supplementary Figure 1C)**.

Also, an AlphaFold2 prediction of Rab29 phosphomimetics showed an “open” conformation around Ser72 **(Supplementary Figure 2A)**, so we decided to assess whether these phosphomimetic Rab29 would be phosphorylated more at Ser72 due to the possibly increased accessibility of this residue. However, no change in phosphorylation of Ser72 was observed either with **(Supplementary Figure 2B)** or without CQ treatment **(Supplementary Figure 2C)**. These data together suggest that phosphorylation at Ser185 does not affect LRRK2 in any way observed.

### PKCα phosphorylates Rab29 at Ser185 and regulates lysosomal localization of Rab29

We next addressed the question as to which kinase would be responsible for this phosphorylation. Kinase determination could be accomplished by *in silico* prediction and following confirmation by *in vitro* experiments. An attempt to predict a kinase for the Ser185 of Rab29 using NetPhos3.1^33^ was made, only to end in no candidates that were scored strong enough, or likely to be a candidate **(Supplementary Table 1)**.

To date, there have been some reports on Rab serine/threonine phosphorylation concerning its C-terminal region. Rab1 and Rab4 is reported to be phosphorylated by CDK1^6^, Rab9 by Ulk1^34^, and Rab11 and Rab37 by PKCα^35,36^ (reviewed in ^7^ and ^4^). Since PKCα was the only hit in the NetPhos3.1 that was also listed as a kinase that phosphorylates Rabs at the C-terminal, we reasoned that this may be a kinase responsible for the phosphorylation of Rab29. Using PKCε as a negative control, we incubated recombinant Rab29 either with PKCα or PKCε *in vitro*. Only Rab29 mixed with PKCα exhibited a band indicative of phosphorylation at Ser185 **(Fig. 5A)**, suggesting PKCα as a kinase for Rab29.

**Figure 5.**
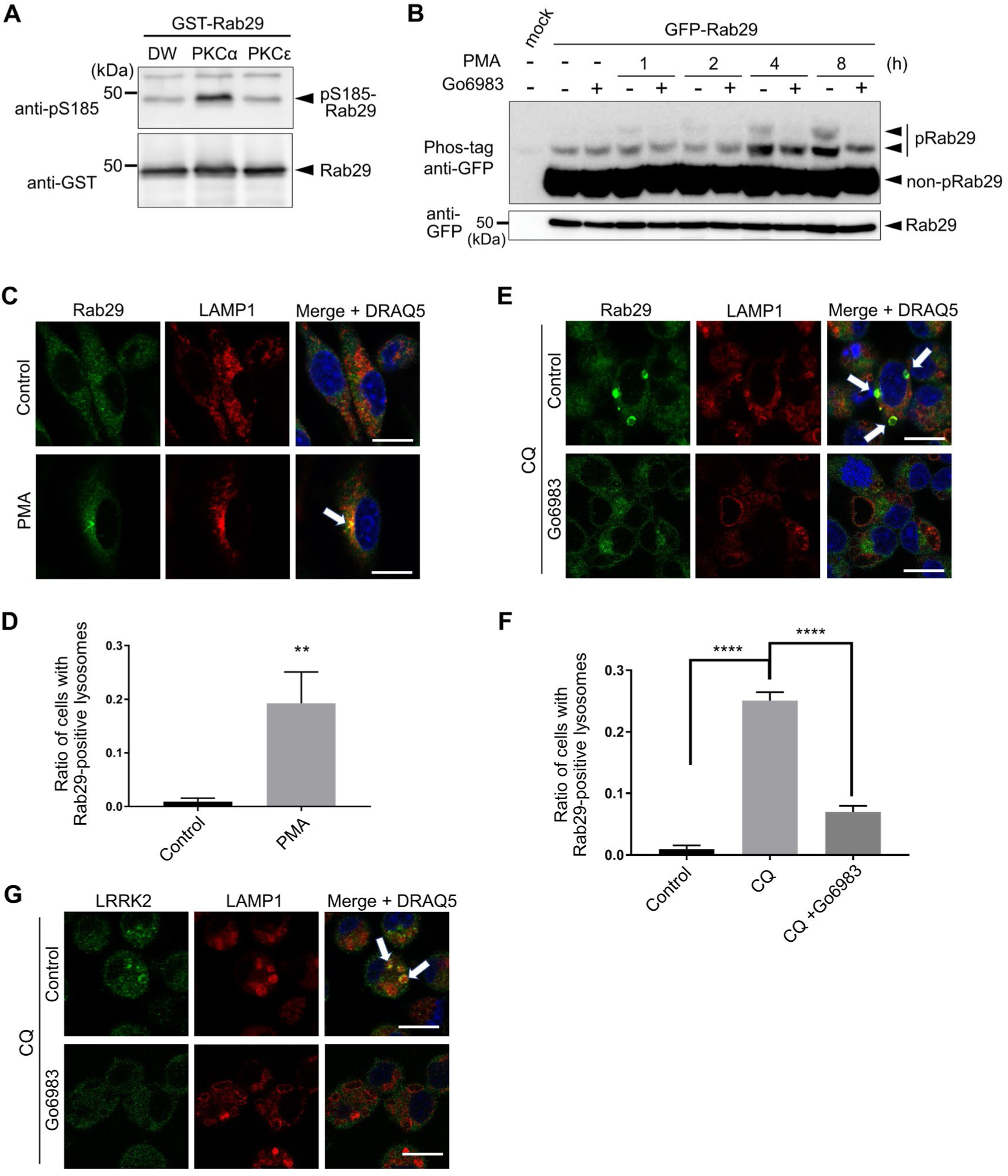
PKCα phosphorylates Rab29 at Ser185 and regulates lysosomal localization of Rab29. **(A)** In vitro kinase assay using recombinant Rab29 and PKCα or PKCε. **(B)** Phosphorylation of Rab29 with PMA or Go6983 over time in HEK293 cells. **(C)** Rab29 localization upon PMA treatment in RAW264.7 cells. The arrow indicates colocalization of endogenous Rab29 with LAMP1. **(D)** Quantification of lysosomal localization of Rab29, as shown in C. Unpaired *t*-test. **: *p* < 0.01. **(E)** Lysosomal localization of endogenous Rab29 upon CQ and Go6983 treatment in RAW264.7 cells. Arrows indicate enlarged lysosomes with Rab29 accumulation. **(F)** Quantification of lysosomal localization of Rab29, as shown in E. One-way ANOVA followed by Tukey’s test. ****: *p* < 0.0001. **(G)** Lysosomal localization of endogenous LRRK2 upon CQ and Go6983 treatment in RAW264.7 cells. Bars = 10 μm.

Phorbol 12-myristate 13-acetate (PMA) is a renowned activator of PKCα. Treatment of HEK293 cells overexpressing Rab29 with PMA resulted in the increase of the levels of phosphorylated Rab29 over time **(Fig. 5B)**. This phosphorylation was canceled upon treatment with a PKC inhibitor Go6983 **(Fig. 5B)**. Thus, we concluded that PKCα is a kinase responsible for phosphorylation of Rab29 in cells.

As changes in Rab29 localization to the lysosomal surface resulted in its phosphorylation **(Fig. 2D)**, we next assessed whether phosphorylation of Rab29 in cell would change its localization. Treatment of RAW264.7 cells with PMA resulted in the translocation of endogenous Rab29 to lysosomes **(Fig. 5C, D)**. In contrast, inhibition of PKCα by Go6983 resulted in a diminution in CQ-induced translocation of endogenous Rab29 to lysosomes **(Fig. 5E, F)**. These data suggest that the PKCα phosphorylation of Rab29 localizes Rab29 to lysosomes. Also considering earlier data that the phosphorylation occurs by forcing the localization of Rab29 to lysosomes, one could infer that Rab29 was stably trapped on lysosomal membranes once phosphorylated. Furthermore, translocation of LRRK2 upon CQ treatment and its inhibition by Go6983 were similarly observed as in the case of Rab29 **(Fig. 5G)**, supporting the link between PKCα and LRRK2 via Rab29.

### LRRK2 is also a regulator of Rab29 localization

To confirm that Rab29 functions upstream of LRRK2, we knocked down the expression of LRRK2 and analyzed the localization of endogenous Rab29 upon CQ treatment. Contrary to expectations, however, knockdown of LRRK2 exhibited lessened accumulation of Rab29 to enlarged lysosomes **(Fig. 6A, B)**. LRRK2 kinase inhibition by MLi-2 also resulted in lower localization of Rab29 to CQ-induced enlarged lysosomes **(Fig. 6C, D)**. These results indicate that, during CQ exposure, LRRK2 is regulated by Rab29, but also acts as a regulator of Rab29 localization in the opposite direction.

**Figure 6.**
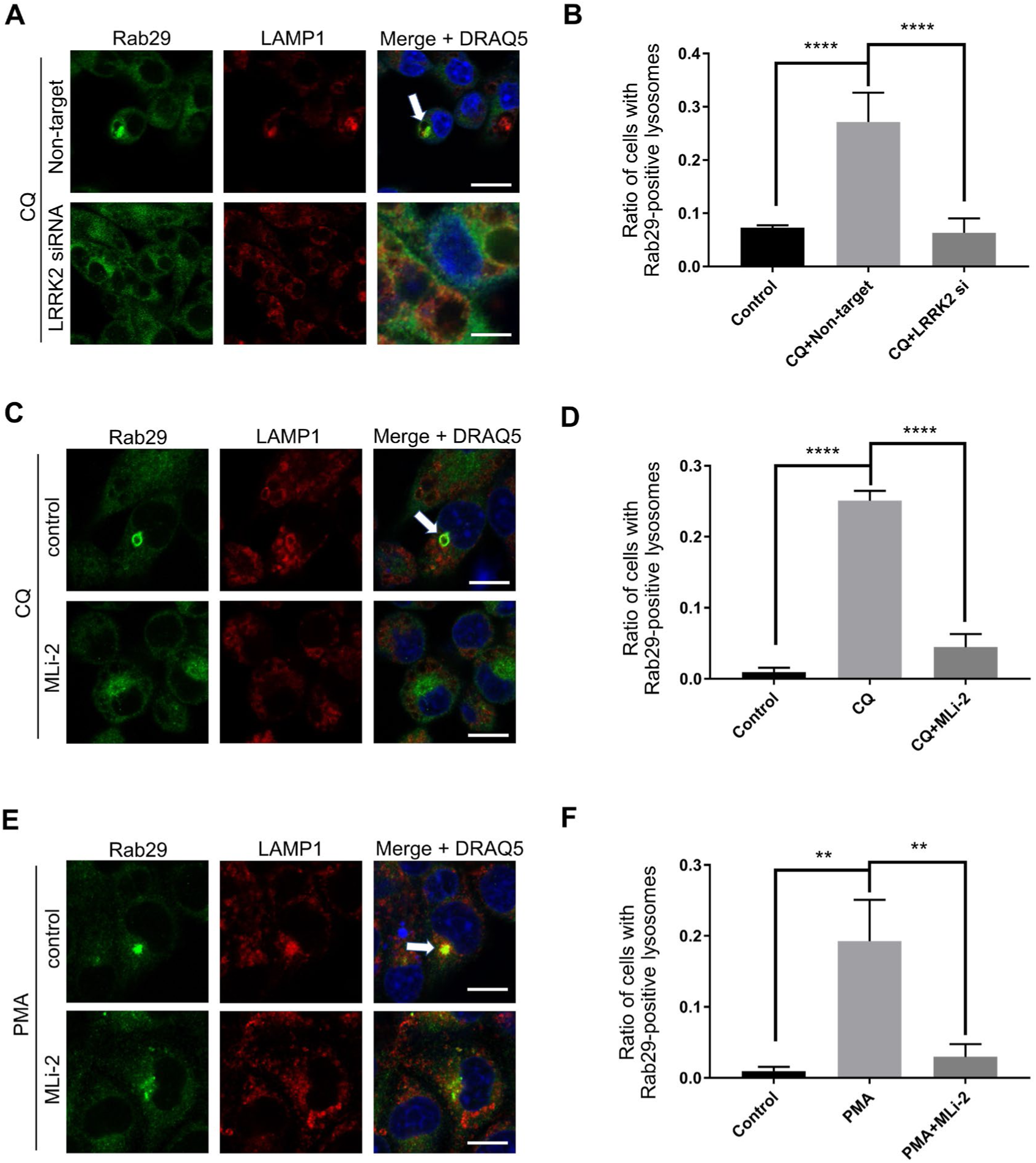
LRRK2 is also a regulator of Rab29 localization. **(A)** Rab29 localization upon knockdown of LRRK2 and CQ treatment in RAW264.7 cells. The arrow indicates enlarged lysosomes with Rab29 accumulation. Bars = 10 μm. **(B)** Quantification of lysosomal localization of Rab29, as shown in A. One-way ANOVA followed by Tukey’s test. ****: *p* < 0.0001. **(C)** Rab29 localization upon MLi-2 (a LRRK2 inhibitor) and CQ treatment in RAW264.7 cells. The arrow indicates enlarged lysosomes with Rab29 accumulation. Bars = 10 μm. **(D)** Quantification of lysosomal localization of Rab29, as shown in C. One-way ANOVA followed by Tukey’s test. ****: *p* < 0.0001. **(E)** Rab29 localization upon MLi-2 and PMA treatment in RAW264.7 cells. The arrow indicates enlarged lysosomes with Rab29 accumulation. Bars = 10 μm. **(F)** Quantification of lysosomal localization of Rab29, as shown in E. One-way ANOVA followed by Tukey’s test. **: *p* < 0.01.

To see whether LRRK2 controls Rab29 localization in other conditions, we turned to PMA treatment which we found would localize Rab29 to perinuclear lysosomes without inducing lysosomal enlargement. Translocation of endogenous Rab29 upon PMA treatment was also suppressed by LRRK2 kinase inhibition, as seen in CQ-treated cells **(Fig. 6E, F)**.

## Discussion

Rab phosphorylation is a common but noteworthy post-translational modification that allows quick regulation and alteration of their functions. However, phosphorylation at the C-terminal region of Rabs is not so common, with only 6 of such described in literature with their pair of kinase^4,7^. Rab11 and Rab37 are reported to be phosphorylated by PKCα, Rab7a by Src, Rab9 by Ulk1, and Rab1 and Rab4 by Cdc2^6,34–37^. Here we provided evidence for another pair, Rab29 and PKCα, which occurs on the lysosomal membrane and could counteract lysosomal overload elicited by a lysosomotropic compound CQ. We also showed that Rab29 localization to lysosomes depends on the phosphorylation by PKCα as well as LRRK2, a known Rab29 interactor. Although the plausible effectors of Rab29 are yet to be discovered, this novel phosphorylation seems to have a function in counteracting lysosomal stress in coordination with LRRK2 and PKCα **(Fig. 7)**.

**Figure 7.**
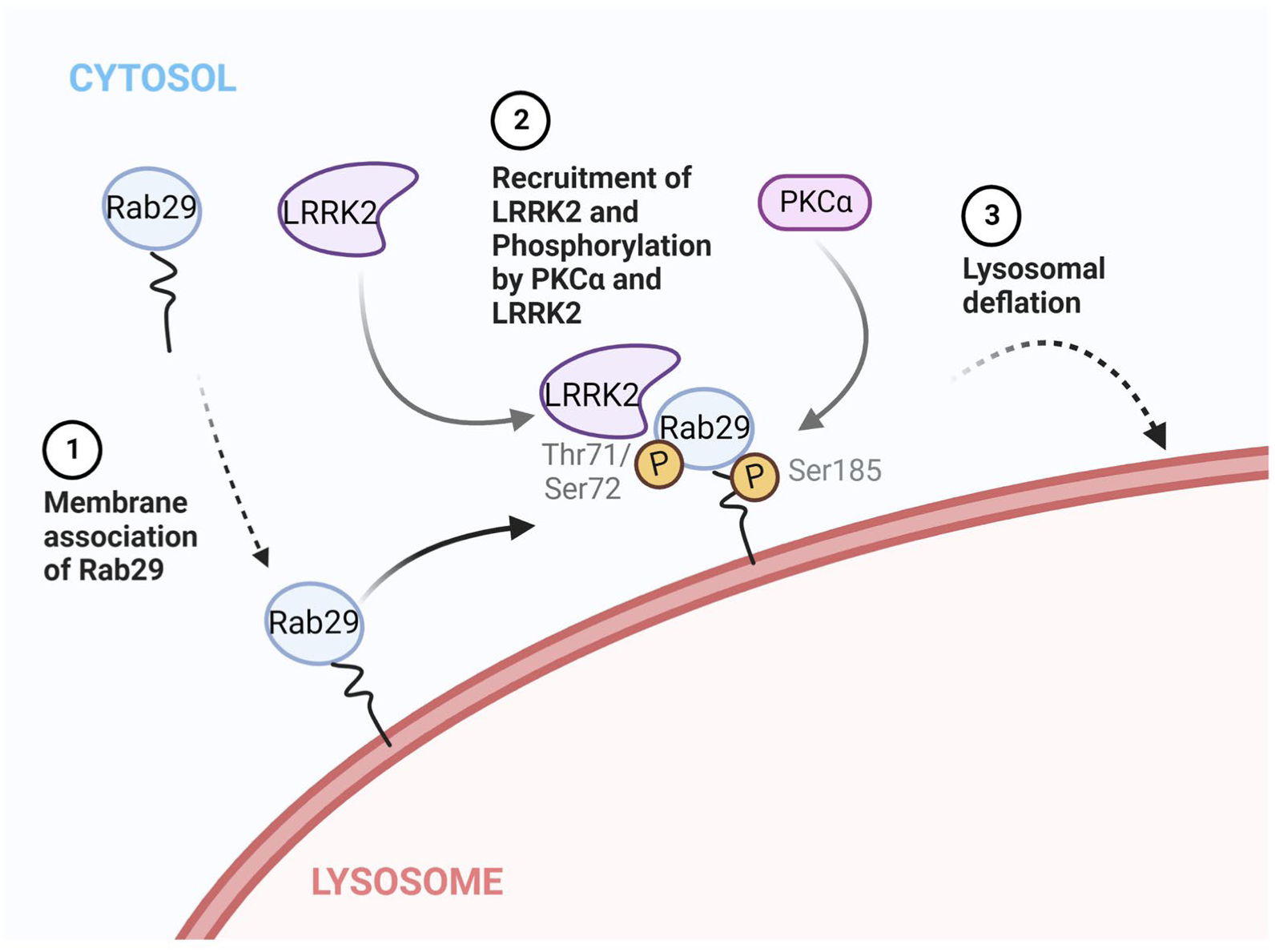
A model for Rab29 translocation, phosphorylation and their effects under lysosomal stress. Upon stimuli that causes lysosomal overload, (1) Rab29 first associates with the lysosomal membranes, and (2) Rab29 recruits LRRK2 to lysosomal membranes and is phosphorylated by PKCα and LRRK2, which stabilizes Rab29 on lysosomal membranes. Then, (3) Rab29/LRRK2 on lysosomes induces downstream effects that lead to lysosomal deflation. This figure was created with BioRender.com.

There have been several reports concerning the relationship between Rab29 and LRRK2, mainly about the phosphorylation and recruitment. Consensus of past literature is that LRRK2 phosphorylates Rab29 (Thr71 and Ser72^10,12^), and Rab29 recruits and activates LRRK2 (either at the Golgi or lysosomes^19,21,38^). The exact mechanism of LRRK2 activation also is unclear, but possibly mediated by heteromultimer formation with Rab29 and/or by membrane association of LRRK2^25,39^. Our data suggest that the regulation of the localization of Rab29 and LRRK2 is intertwined, both needing each other to be stably localized on membranes. A recent report showed that LRRK2 has two different sites to which Rabs can bind, one for unphosphorylated Rabs, and one exclusively for phosphorylated Rab8 or Rab10^26^. Inhibition of LRRK2 kinase activity showed dispersed Rab29 localization (Fig. 6C, D), so it could be reasoned that the binding of LRRK2-phosphorylated Rab29 to LRRK2 at multiple sites may be necessary for its activation or stabilization at the lysosomal membrane.

PKCα is a ubiquitously expressed kinase implicated in a multitude of pathways from cell proliferation to apoptosis^40^. PKCα is mainly localized at the plasma membrane in its active state but could be transported via the endosomal pathway while keeping its active state upon PMA treatment^41^. So, the perinuclear localization of Rab29 that overlapped with LAMP1 upon PMA treatment (Fig. 5C) could be explained in this context, where active PKCα happened to encounter Rab29 somewhere on the endosomal pathway and the resulting phosphorylation caused the change in its localization.

The exact mechanism of Rab29 localization to LAMP1-positive compartments, especially how Rab29 gets ‘trapped’ on the membrane where it is or has been phosphorylated, is still unclear. Usually, Rab localization is strictly regulated by their GEFs, and further investigation is needed to answer whether this phenomenon is also due to GEF and its activity. Recently, a GEF of Rab29 was proposed to be Rabaptin5^42^, and it might be worthwhile to assess their interactions in similar conditions. There might also exist unknown interactors specific to Rab29 phosphorylated at Ser185 that mediate Rab29 localization.

We have previously reported that the knockdown of Rab29 causes enhanced enlargement of lysosomes and impaired release of lysosomal contents upon exposure to CQ^19^. The amelioration of lysosomal enlargement with phosphomimetic Rab29 overexpression (Fig. 4B, C) could be explained as the reverse of knockdowns, and lysosomal content release downstream of LRRK2 could have had an effect on regulating the size of lysosomes. We were not able to assess this possibility due to technical difficulties concerning cell types.

Recently, inhibition of PKC has been reported to prevent aggregation of α-synuclein, a key causative protein of PD, upon transfection with pre-aggregated “seed” α-synuclein^43^. In our data, PKC inhibition resulted in dispersed Rab29 localization under lysosomal stress (Fig. 5E). Considering that most pathogenic LRRK2 mutations lead to increased phosphorylation of its substrates, either by upregulation of kinase activity or enhanced substrate binding, and that Rab29 activates LRRK2, our data is in line with this view and may further link the abnormalities in Rab29 with PD in terms of pathological mechanisms. This also highlights the importance of analyzing the detailed functions of Rab29 at lysosomes where it may respond to intracellular stress.

In summary, our data provide evidence for a novel phosphorylation in Rab29 by PKCα capable of controlling Rab29 localization in concert with LRRK2. These phosphorylations and the following localization change were considered to be important for maintaining lysosomal morphology upon lysosomal overload stress. Further studies would be needed to clarify how this stress response mechanism is initiated, whether it is regulated by other key molecules, and whether it could be involved in the pathogenesis of PD.

## Materials and Methods

### Antibodies

The following antibodies were used in this study: GFP (Invitrogen, A11122), LAMP1 (BD Pharmingen, H4A3), phospho-Ser72-Rab29 (generated in our previous study^11^), FLAG (MBL, FLA-1), LRRK2 (MJFF, MJFF2), α-tubulin (Abcam, DM1A), non-phospho-Ser185-Rab29 (generated in this study), phospho-Ser185-Rab29 (generated in this study), LAMP1 (Bio-Rad, 1D4B), Rab10 phospho-Thr73 (Abcam, MJF-R21), Rab10 (Cell Signaling Tech, D36C4), Rab29 (Abcam, MJF-R30-124), GST (GE Healthcare, 27-4577-01), and secondary antibodies. Secondary antibodies for immunocytochemistry were goat/donkey anti-IgGs labeled with either Alexa-488, Alexa-546, Alexa-555 or Alexa-647 (Thermo Fisher Scientific). Secondary antibodies for immunoblotting were HRP-labeled anti-IgGs (Jackson Immunoresearch).

### Reagents

The following reagents were used at final concentrations as indicated: Chloroquine (CQ) (50 μM, Sigma Aldrich), phorbol 12-myristate 13-acetate (PMA) (100 nM, Santa Cruz), MLi-2 (50 nM, Abcam), Go6983 (200 nM, Sigma Aldrich), AP21967 (1 μM, TaKaRa).

### Generation of phospho-Ser185-Rab29 specific antibodies

To generate human phospho-Ser185-Rab29 specific antibody, rabbits were immunized with KLH-conjugated peptides (KLH-RNSTEDIMSL(pS)TQGD, KLH-RNSTEDIMSLSTQGD, human Rab29 sequences around Ser185) 3 times with a 2 week’s interval. Serum was collected 6 weeks after the final immunization, and purified by dual affinity purification, which is a method using non-phosphorylated peptides as a first column and purifying the flow-through with phosphorylated peptides, ensuring more affinity to phosphorylated peptides. These processes were performed at Biologica Co.

### Plasmids and siRNA

The plasmid pEGFP-C1-human Rab29 was generated by inserting human Rab29 sequence from pFN21A-Halo-Rab29 (Promega, #FHC08084) into Bgl II - Eco RI site of pEGFP-C1-rat Rab29 plasmid that was used previously^11^. Phospho-mutants of Rab29 were generated by a site-directed PCR mutagenesis protocol. A set of plasmids encoding

EGFP-mouse Rabs was provided by Dr. Mitsunori Fukuda (Tohoku Universty). LAMP1-FRB and FRB-Fis1 plasmids were provided by Dr. Richard J. Youle (NIH)^44^. The plasmid encoding 2×FKBP-GFP-Rab29 was generated by transferring 2×FKBP sequence from Addgene plasmid #20149 into Hind III site of pCMV10 plasmid followed by inserting EGFP-Rab29 sequence into Not I - Xho I site of pCMV10. The siRNAs used were purchased from Dharmacon (siGenome smart pool), Thermo Fisher Scientific, and Bioneer. The siGenome smart pool is a cocktail of 4 different oligonucleotides to reduce off-target effects.

### Cell culture and transfection

Human embryonic kidney cell line HEK293 and mouse macrophage-like cell line RAW264.7 (purchased from ECACC) were cultured in DMEM (Dulbecco’s modified Eagle’s medium, Sigma Aldrich) with 10% FBS (Fetal bovine serum, Biowest) and 1% PS (Penicillin/Streptomycin, Gibco) under 5% CO_2_, 37°C. RAW264.7 cells were activated by IFN-γ (15 ng/mL) 48 hours prior to analysis. HEK293 and RAW264.7 cells were cultured on Tissue Culture-treated culture dish (Corning) and Petri dish for suspension cell culture (Sumitomo Bakelite), respectively. Passage of each cell line was done by pipetting off attached cells.

Transfection of plasmid vectors to HEK293 cells was conducted using Lipofectamine 2000 (Thermo Fisher Scientific) and Thermo Fisher Scientific’s recommended protocols for transfection using Lipofectamine 2000. Transfection of siRNA to RAW264.7 cells was conducted using Lipofectamine RNAiMAX (Thermo Fisher Scientific) and Thermo Fisher Scientific’s recommended protocols for reverse transfection.

### Treatment of reagents to cells

To HEK293 cells, each reagent was treated for 24 hours. To RAW264.7 cells, each reagent was treated for 3 hours. The final concentrations used were described in Reagents section unless otherwise stated.

### Cell culture and fixation for Immunocytochemistry

Cells intended for ICC were cultured on cover glasses (Matsunami) coated with poly-D-lysine (PDL). Coating was done by incubating 200 μL of PDL, diluted to 50 μg/mL by DPBS, on each cover glass for more than 30 min at 37°C, then washed several times with DPBS. Cover glasses were washed 2 times with 1M NaOH for an hour each, then washed with 70% ethanol for 2 times or more and stored in 70% ethanol at 4°C.

Fixing of cells was conducted by immersing cover glasses in 4% PFA in DPBS for 30 minutes at room temperature. After fixation, cover glasses were washed with DPBS and immersed in 100% ethanol for sample dehydration at -20°C. Samples were stored in this state until further means of immunocytochemistry.

### Immunocytochemistry

Samples stored in 100% ethanol were washed with DPBS before blocking with blocking buffer (3% (w/v) BSA, 0.1∼0.5% Triton X-100 in DPBS) for 30 minutes at room temperature. Samples were then incubated with primary antibodies diluted in blocking buffer for 3 hours at room temperature or overnight at 4°C. Each cover glass was placed on a 35 μL spot of diluted primary antibody solution made on a sheet of parafilm (Bemis) so that the surface with cells present is facing the antibody solution^45^. After washing with DPBS 3 times for 5 minutes, samples were placed on secondary antibodies (1:500 dilution for Alexa conjugated anti-IgG, 1:2000 for DRAQ5) for 1 hour at room temperature or overnight at 4°C. Followed by further washes by DPBS for 3 times, samples were mounted on slide glasses (Matsunami) with 7 μL of Permafluor mountant (Thermo Fisher Scientific) for each cover glass.

### Preparation of cell lysates

Cultured cells were washed with DPBS, then scraped off in lysis buffer (50 mM Tris-HCl, pH = 7.6, 150 mM NaCl, 0.5% Triton X-100, 0.3 tablets of cOmplete protease inhibitor cocktail, 2 tablets of Phos-STOP phosphatase inhibitor cocktail) using a micropipette tip whose tip has been cut off. The scraped cell-lysis buffer solutions were rotated in a 1.5 mL tube at 4°C for 30 minutes, centrifuged at 17,300 × *g*, 4°C and the supernatant was collected as cell lysate, either stored at -20°C or -80°C, or proceeded on to each experiment.

### Immunoprecipitation

Immunoprecipitation was performed using either Protein G agarose (Invitrogen) or GFP-Trap beads (Chromotek). For immunoprecipitation using Protein G agarose, cells were first precleared of nonspecific agarose or protein G binding by rotating with washed Protein G agarose for 30 minutes at 4°C. Part of the supernatant was collected as the input fraction, and the rest was mixed with washed Protein G agarose and 1 μL of antibody per 500 μL of cell lysate and rotated for 3 hours at 4°C. Agarose beads were then washed with TBS containing cOmplete and Phos-STOP for 3 times and boiled in 1.5× sample buffer for 10 minutes at 90°C.

For immunoprecipitation using GFP-Trap beads, cell lysates were first diluted with TBS buffer (50 mM Tris-HCl, pH = 7.6, 150 mM NaCl) to reduce the concentration of Triton X-100. These were next mixed with equilibrated and washed GFP-Trap beads, then rotated for 2 hours at 4°C. The samples were then centrifuged at 2500 × *g* to separate the unbound supernatant from the beads. The beads were washed with TBS buffer for 3 more times, and were boiled in 2× sample buffer for 10 minutes at 90°C.

### In vitro kinase assay

Previously purified Recombinant Rab29^11^ (450 ng) and PKCα or PKCε (Aviva Systems Biology) (50 ng each) were mixed in kinase assay buffer (50 mM Tris-HCl, 10 mM MgCl_2_, 1 mM CaCl_2_, 2 mM DTT, 333 nM PMA, 1 mM ATP) and incubated at 30°C for 30 minutes on a shaker at 900 rpm. The mixtures were then diluted in 4× sample buffer (4×LDS Buffer mixed with 4% (v/v) 2-mercaptoethanol) and boiled at 90°C for 5 minutes before proceeding on to SDS-PAGE.

### SDS-PAGE and Western blotting

For samples that need quantification, BCA assays using BCA assay kit (TaKaRa) were performed before boiling the lysed cells or medium mixed with 1/3 volume of 4× sample buffer for 10-15 minutes at 90°C. SDS-PAGE was conducted to separate proteins by their size using either 7.5%, 10% or 12.5% Tris-glycine gels. Phos-tag SDS-PAGE was performed to separate phosphorylated proteins from non-phosphorylated forms using Phos-tag gels (7.5% Tris-glycine gels supplemented with 150 μM MnCl_2_ and 75 μM Phos-tag acrylamide (Wako)). For each gel, 1 lane was kept for molecular weight markers (Precision Plus Dual Color Protein Standard, Bio-Rad). Samples after electrophoresis were transferred to PVDF membranes (Millipore) in blotting buffer (10 or 20% methanol, 25 mM Tris-HCl, 200 mM glycine). Phos-tag gels were rocked in blotting buffer with EDTA to remove excess Mn^2+^ ions. PVDF membranes were then blocked in 5% skim milk in TBS-Tween (Tris-buffered saline (50 mM Tris, 150 mM NaCl, pH = 7.6) mixed with 0.1% Tween 20 (Sigma Aldrich)), or for samples subjected to phospho-antibodies, 5% bovine serum albumin (Sigma Aldrich) in TBS-Tween, for 30 minutes at room temperature. Primary antibodies were diluted in Immuno-enhancer (Wako) and incubated with blocked PVDF membranes overnight at 4°C. After subsequent washing with TBS-Tween 3 times for 5 minutes each, membranes were incubated with secondary antibodies diluted in Immuno-enhancer for 45 minutes at room temperature or overnight at 4°C. Then, membranes were washed 2 times for 10 minutes each before being immersed in Immunostar reagents (Wako) for chemiluminescence. Chemiluminescent signals were detected by LAS-4000 mini (FUJIFILM) and quantified using Fiji^46^.

### Mass spectrometry

For identifying the phosphorylation site of Rab29, the part of the gel corresponding to phosphorylated Rab29 was cut after Phos-tag SDS-PAGE and digested by either trypsin or a combination of chymotrypsin and elastase using a ProGest robot (DigiLab). The digested samples were then loaded on a nano LC-MS/MS with a Waters NanoAcquity HPLC system interfaced to a ThermoFisher Q Exactive. The retrieved data were then analyzed using a local copy of Mascot (Matrix Science). The procedures after cutting the gel were performed at Filgen Inc.

### Quantification of lysosomal size

Lysosomal cross-sectional areas in images obtained from ICC were quantified using Fiji^46^. Image acquisition was performed at *z* levels where lysosomes would be the largest. Lysosomes were traced with an oval tool and their surface areas were obtained by the measure command. The largest lysosome of all visible cells in the images were traced and measured. The surface areas were then recorded in a Microsoft Excel sheet for further analysis.

### Prediction of kinases

The full-length sequence of human Rab29 was analyzed by NetPhos3.1^33^.

### Molecular modeling

The structural modeling of proteins was conducted using UCSF ChimeraX^47^, developed by the Resource for Biocomputing, Visualization, and Informatics at the University of California, San Francisco, with support from National Institutes of Health R01-GM129325 and the Office of Cyber Infrastructure and Computational Biology, National Institute of Allergy and Infectious Diseases.

### Statistical analyses

Statistical analyses were conducted using R (The R Foundation) or GraphPad Prism 7. For multiple comparisons after ANOVA analysis, Tukey’s test was performed when comparing every mean with every other mean, and Dunnett’s test was performed when comparing every mean with a control mean. Statistical significance was set to *p* < 0.05.

### Data availability

All data sets used or analyzed in this study are available from the corresponding author upon request.

## Acknowledgements

We thank Dr. Richard J. Youle (NIH) for providing us with plasmids encoding FRB, Dr. Mitsunori Fukuda for a set of GFP-Rab constructs, and our lab members for helpful suggestions and discussions. The study was supported by JSPS KAKENHI grant numbers 16K07039 (T. Kuwahara), 19K07816 (T. Kuwahara), 20H00525 (T. Iwatsubo), 20J12819 (T. Komori) and 21J12881 (M. Sakurai).

## Author contributions

T. Komori, T. Kuwahara and T. Fujimoto conceived and designed studies. T. Komori, T. Kuwahara, T. Fujimoto and M. Sakurai preformed the experiments. T. Komori, T. Kuwahara and T. Iwatsubo wrote the manuscript. All authors read and approved the final manuscript.

---

**Figure S1.**
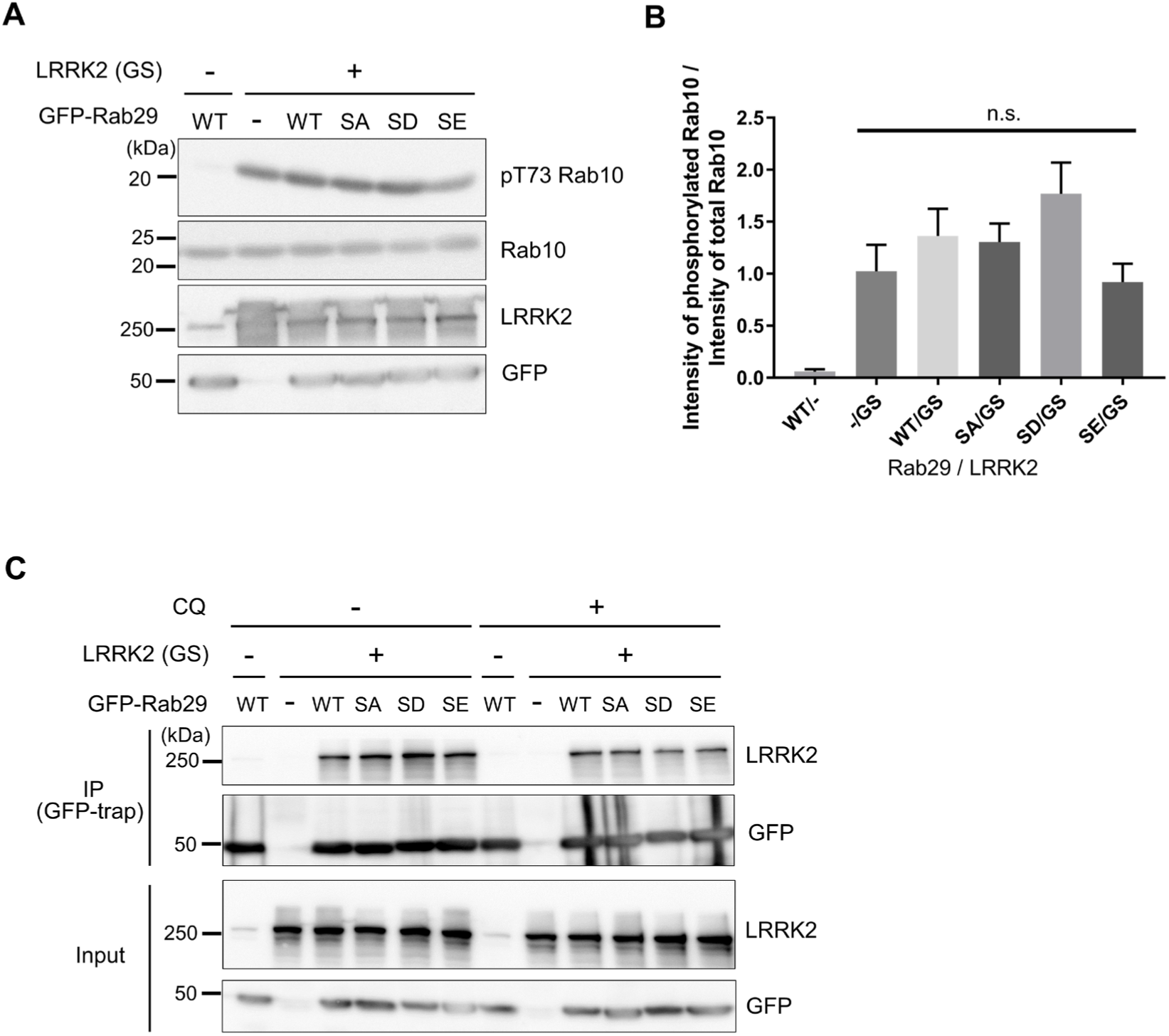
Rab29 Ser185 phosphomimetics do not alter LRRK2 kinase activity or Rab29-LRRK2 interaction. **(A)** Phosphorylation of Rab10 by LRRK2 in HEK293 cells upon overexpression of Rab29 wild-type (WT), S185A mutant (SA) or S185D/E phosphomimetics (SD, SE). GS: G2019S mutant. **(B)** Quantitative analysis of Rab10 phosphorylation, as shown in A. One-way ANOVA followed by Dunnett’s test against control (WT Rab29/LRRK2 G2019S expression). n.s.: not significant. **(C)** Co-immunoprecipitation of G2019S LRRK2 by GFP-Rab29 in HEK293 cell lysates using an anti-GFP antibody (GFP-trap).

**Figure S2.**
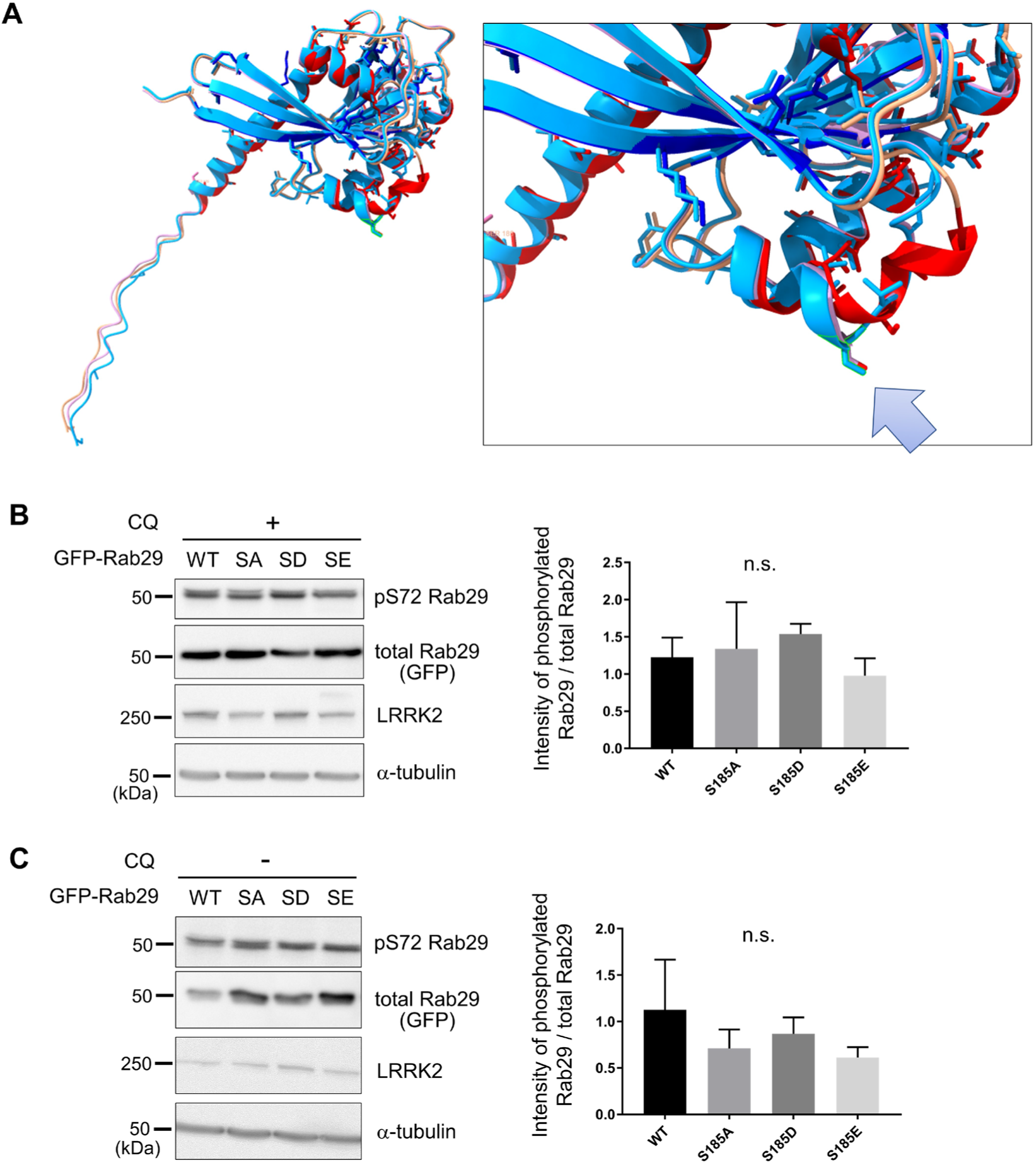
Rab29 Ser185 phosphomimetics do not alter their phosphorylation by LRRK2 at Ser72. **(A)** An AlphaFold2 prediction of S185D and S185E phosphomimetics of Rab29 (colored light blue and pink, respectively) and an inset around the switch II region. The arrow indicates Ser72 residue. **(B)** Phosphorylation of Rab29 at Ser72 in HEK293 cells overexpressing Rab29 S185A mutant (SA) or S185D/E phosphomimetics (SD, SE) upon CQ treatment. One-way ANOVA followed by Dunnett’s test against WT. n.s.: not significant. **(C)** Phosphorylation of Rab29 at Ser72 in HEK293 cells overexpressing Rab29 S185A mutant (SA) or S185D/E phosphomimetics (SD, SE) without CQ treatment. One-way ANOVA followed by Dunnett’s test against WT. n.s.: not significant.

**Table S1.** Prediction of kinases for Rab29 Ser185 by NetPhos 3.1. The full-length sequence of human Rab29 was analyzed by NetPhos3.1^33^. The candidate kinases that could phosphorylate Rab29 at Ser185 are shown in order of likelihood. All of them have low scores below 0.5, meaning that they are unlikely to be the responsible kinase for Ser185.

| Sequence | # | x | Context | Score | Kinase | Answer |
| --- | --- | --- | --- | --- | --- | --- |
| sp_O14966_RAB7L_HUMA | 185 | S | IMSLSTQGD | 0.478 | CKII | . |
| sp_O14966_RAB7L_HUMA | 185 | S | IMSLSTQGD | 0.456 | cdc2 | . |
| sp_O14966_RAB7L_HUMA | 185 | S | IMSLSTQGD | 0.45 | GSK3 | . |
| sp_O14966_RAB7L_HUMA | 185 | S | IMSLSTQGD | 0.412 | CaM-II | . |
| sp_O14966_RAB7L_HUMA | 185 | S | IMSLSTQGD | 0.368 | CKI | . |
| sp_O14966_RAB7L_HUMA | 185 | S | IMSLSTQGD | 0.355 | DNAPK | . |
| sp_O14966_RAB7L_HUMA | 185 | S | IMSLSTQGD | 0.355 | ATM | . |
| sp_O14966_RAB7L_HUMA | 185 | S | IMSLSTQGD | 0.275 | p38MAPK | . |
| sp_O14966_RAB7L_HUMA | 185 | S | IMSLSTQGD | 0.268 | RSK | . |
| sp_O14966_RAB7L_HUMA | 185 | S | IMSLSTQGD | 0.259 | PKG | . |
| sp_O14966_RAB7L_HUMA | 185 | S | IMSLSTQGD | 0.222 | PKA | . |
| sp_O14966_RAB7L_HUMA | 185 | S | IMSLSTQGD | 0.203 | PKC | . |
| sp_O14966_RAB7L_HUMA | 185 | S | IMSLSTQGD | 0.167 | cdk5 | . |
| sp_O14966_RAB7L_HUMA | 185 | S | IMSLSTQGD | 0.085 | PKB | . |
| sp_O14966_RAB7L_HUMA | 185 | S | IMSLSTQGD | 0.075 | unsp | . |

